# GABA Increases Sensory Transmission In Monkeys

**DOI:** 10.1101/2023.12.28.573467

**Authors:** Amr A. Mahrous, Lucy Liang, Josep-Maria Balaguer, Jonathan C. Ho, Krishnapriya Hari, Erinn M. Grigsby, Vahagn Karapetyan, Arianna Damiani, Daryl P. Fields, Jorge A. Gonzalez-Martinez, Peter C. Gerszten, David J. Bennett, C.J. Heckman, Elvira Pirondini, Marco Capogrosso

## Abstract

Sensory input flow is central to voluntary movements. For almost a century, GABA was believed to modulate this flow by inhibiting sensory axons in the spinal cord to sculpt neural inputs into skilled motor output. Instead, here we show that GABA can also facilitate sensory transmission in monkeys and consequently increase spinal and cortical neural responses to sensory inputs challenging our understanding of generation and perception of movement.

## MAIN TEXT

Movement perception is encoded by an ensemble of neural elements that continuously convey sensory information on limb position and velocities to the central nervous system. This information flow not only has a perceptual value in our conscious experience of movement, but also plays a major role in shaping neural activity of spinal motoneurons. Glutamatergic inputs from proprioceptive Ia-afferents to motoneurons are one of the strongest excitatory pathways in the nervous system [1, 2]. The potency of this monosynaptic circuit is such that, if left unmodulated, it could overwhelm spinal motoneurons with thousands of excitatory inputs per second rendering fine motor skills impossible [3]. For this reason, it is believed that a presynaptic modulation mechanism capable of gating and filtering this input would be necessary to enable volitional neural control of spinal motoneurons [3, 4]. In fact, almost 130 years ago primary afferent depolarization (PAD), an inward current in sensory axons, was observed upon electrical stimulation of nerves [5]. The co-occurrence of PAD with instances of reduced monosynaptic responses in motoneurons led to the concept of presynaptic inhibition: a mechanism that limits neurotransmission from afferents to motoneurons. PAD was thought to be the cause and electrophysiological signature of presynaptic inhibition [4, 6, 7]. Since early experiments it was clear that PAD had two unique features: first, it is a purely axonic mechanism mediated by GABAergic axo-axonic synapses onto sensory axons [3, 8, 9] (**Fig. 1A**); second, although mediated by GABA_A_ receptors [3, 8, 10], PAD is a depolarization event because sensory afferents have a depolarized Cl^-^ equilibrium potential due to expression of NKCC1 transporter and lack of KCC2, making GABA excitatory [11–13] (**Fig. 1A**). These facts generated an apparent contradiction in the link between PAD and presynaptic inhibition: PAD is excitatory at face-value but causes inhibition. To solve this dilemma, it was hypothesized that excitatory GABA_A_-generated currents shunted afferent terminals thereby preventing further glutamate release onto motoneurons [14–16]. However, recent works showed that GABA_A_ receptors are not expressed at Ia-afferent terminals but rather at nodes and branch points [17, 18], and suggested that the association between PAD and presynaptic inhibition was a correlation rather than a causal link [18–23]. Since modulation of sensory inputs is thought to be critical to skilled motor control [24, 25], here we tackled this controversy directly in monkeys, the most relevant animal model for skilled motor control. Although some studies in monkeys have evaluated presynaptic inhibition of reflexes [26, 27], none has directly measured the classic long-lasting PAD commonly observed in other species [3, 18, 28, 29] to establish causal relationships.

**Fig. 1.**
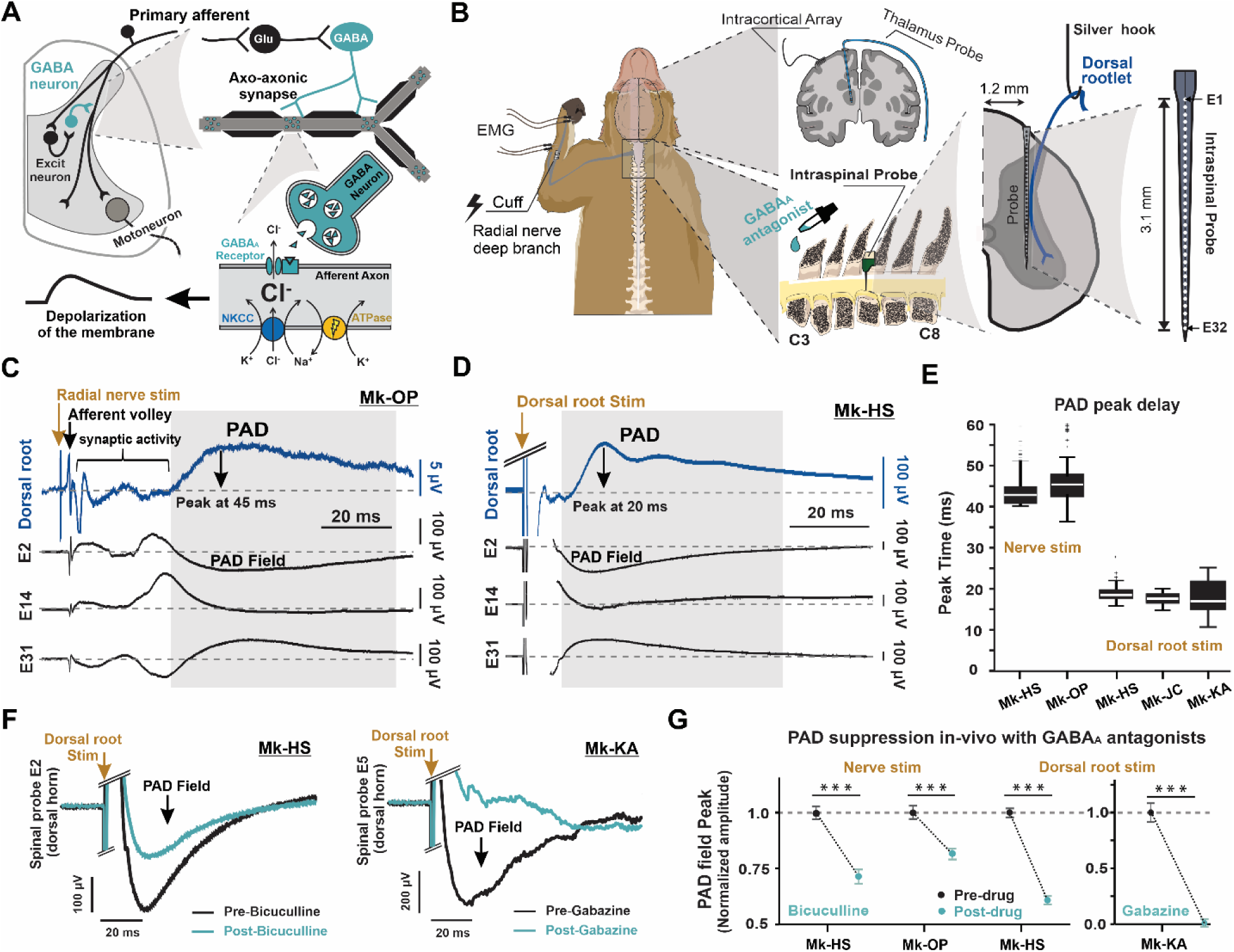
Primary afferent depolarization (PAD) in non-human primates is mediated by GABA_A_ receptors. **A:** Diagram of the tri-synaptic PAD circuit in the spinal cord. An excitatory neuron activated by primary afferents excites a GABA interneuron which forms axo-axonic synapses with primary afferents at nodes and branching points. Unlike other neurons, sensory afferents maintain a high concentration of intracellular chloride ions via expression of the NKCC1 transporter. When GABA binds to GABA_A_ receptors on the axon, chloride ions leave the axon causing depolarization (primary afferent depolarization, PAD). **B:** Schematic representation of the in-vivo experimental setup in anesthetized monkeys. PAD was recorded from cervical dorsal roots using a silver hook electrode, and motor output was recorded as the EMG of hand and wrist muscles using needle electrodes. Intraspinal and supraspinal neuronal activity was recorded through multi-electrode arrays implanted in spinal cervical segments, the thalamus, and sensory cortex. The diagram on the right shows the mediolateral location and depth of the 32-channels spinal linear array. **C-D:** Classic PAD signal recorded from a C8 dorsal rootlet (blue) along with PAD field across multiple spinal laminae (black traces) recorded simultaneously using the linear array inserted at C6. PAD was evoked either by a single pulse stimulus to the ipsilateral radial nerve (C, 0.1 ms pulse) or C6 dorsal root (D, 0.4 ms pulse). Note that the PAD field in the spinal cord is largest in the most dorsal electrode contacts, minimal around the middle, and reverses polarity at the most ventral contacts. **E:** Delay of the peak response of PAD in the dorsal root for nerve stimulation (2 box plots on the left) and dorsal root stim (3 box plots on the right) from multiple experiments. For all boxplots, the whiskers extend to the maximum and minimum, excluding outliers. Central line, top, and bottom of box represent median, 75th, and 25th percentile, respectively. **F:** PAD field recorded in the dorsal aspect of the cord using the array in response to a single stimulus to the dorsal root before (black) and after (cyan) intrathecal administration of GABA_A_ antagonists (left: bicuculline, right: gabazine) in different monkeys. **G:** Mean and standard error plot of PAD field peak amplitude (normalized to the mean before drug administration) evoked by radial nerve or dorsal root stimulation, showing reduction of PAD after bicuculline or gabazine administration (***: P<0.001; two-tailed bootstrapping was used for pre and post drug conditions, with 258 and 121 points for Mk-HS nerve stim, 185 and 189 points for Mk-OP nerve stim, 218 and 201 points for Mk-HS DR stim, 107 and 84 points for Mk-KA DR stim).

We engineered a high throughput set-up in anesthetized monkeys (**Fig. 1B**) enabling manipulation of PAD and recording of its downstream effects at multiple locations along the nervous system. To evoke PAD, we stimulated the sensory afferents at motor threshold through a cuff electrode on the deep branch of the radial nerve (purely proprioceptive, no cutaneous afferents) or through dorsal root stimulation at the cervical segments. Using a silver hook electrode, we measured PAD in sensory afferents from the proximal portion of a dorsal rootlet that was severed close to its entry zone (**Fig. 1B**). With this technique, we could observe changes in intra axonal potentials with high-fidelity [8, 18, 28]. We consistently observed an afferent volley (2-3 ms after nerve stimulation) and a synaptic field followed by clear PAD that peaked around 15-20 ms for post-dorsal root stimulation (**Fig. 1D and 1E**) and 40-45 ms post-nerve stimulation (**Fig. 1C and 1E**) and lasting 50 - 100 ms. We validated our observations by inspecting extracellular fields recorded by a high-density dorso-ventral linear probe at the C6-C7 spinal segments (**Fig. 1B**). In the gray matter, PAD fields clearly appeared throughout different spinal laminae (opposite polarity in the ventral horn) and matched the time course of PAD in the dorsal rootlet (**Fig. 1C-D** and **Extended Data Fig 1**).

We then tested whether PAD is mediated by GABA_A_ receptors, as in other species. For this, we applied a GABA_A_ antagonist, either bicuculline or gabazine, intrathecally. Both drugs significantly reduced the amplitude of intra-spinal PAD fields (**Fig. 1F-G**) and dorsal root PAD (**Extended Data Fig. 2A-B**) for both types of stimuli, indicating that GABA_A_ plays a major role in generation of PAD in primates. Importantly, the reduction in PAD was not caused by changes in the afferent volley entering the spinal cord (**Extended Data Fig. 2C-D**). As expected from GABA blockers, both drugs caused an overall increase of background neural activity (**Extended Data Fig. 3**), likely in consequence of reduced GABA-mediated inhibition in the grey matter and therefore a general increase in excitability.

**Fig. 2.**
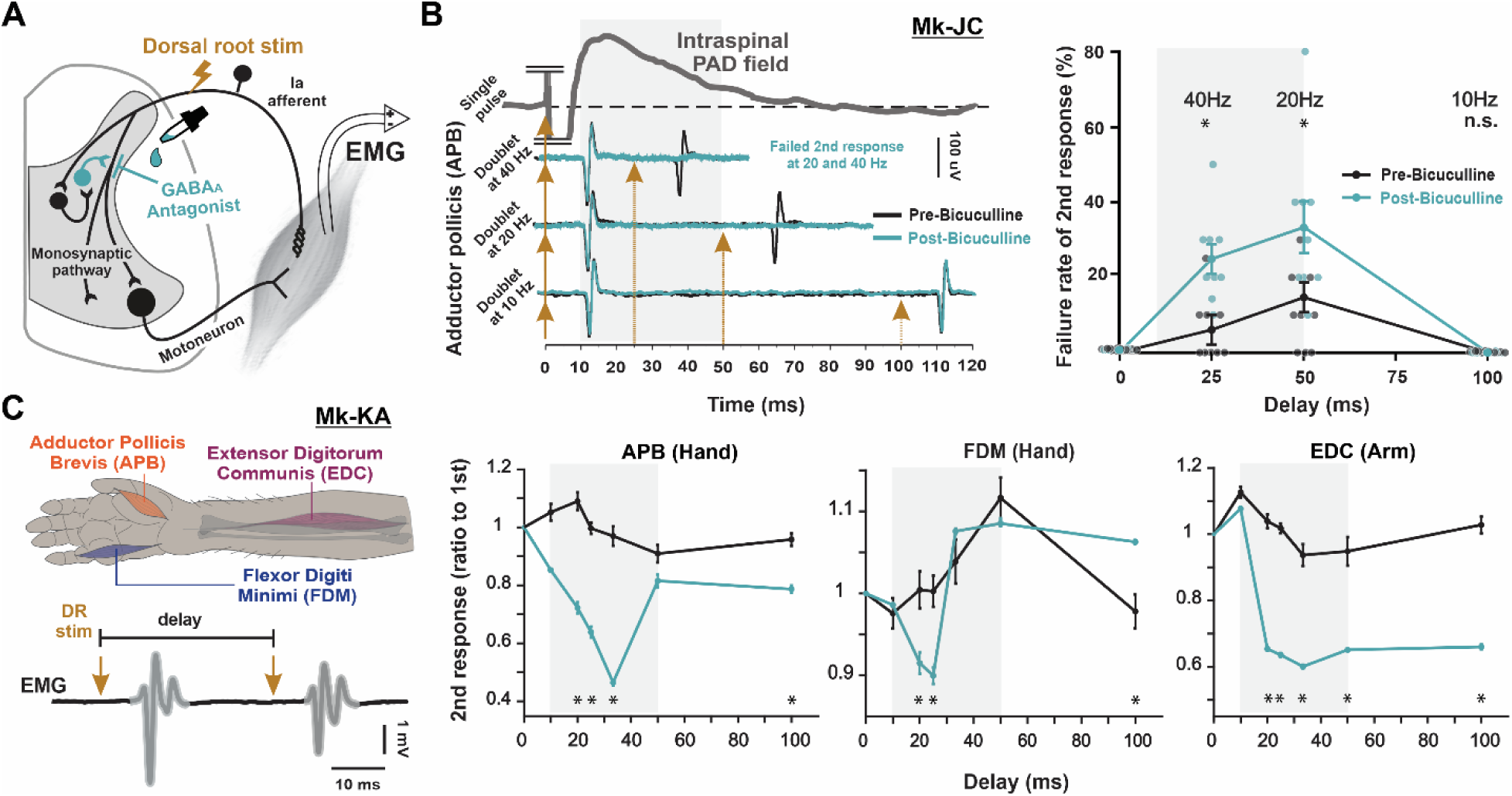
Motor responses are suppressed after blocking GABA_A_ receptors. **A:** Schematic diagram of primary afferent monosynaptic pathway to motoneurons along with PAD circuit. **B: Left:** top trace (grey) shows the time course of PAD field in the dorsal column (recorded using the intraspinal probe) in response to a single C8 dorsal root stimulus in Mk-JC. Lower traces show, in the same animal, EMG responses of the APB muscle to dorsal root stimulation at the motor threshold with two successive pulses (doublets) at 40, 20, and 10 Hz before (black) and after (cyan) intrathecal administration of bicuculline. **Right:** Mean and standard error plot shows the increased failure rate of the second response in a doublet at 20 and 40 Hz, but not at 10 Hz after bicuculline administration. The response to the first pulse of all doublets did not show any failure before or after drug administration. *: P<0.001; two-tailed bootstrapping with 9 points for each delay, and each pre and post drug condition. **C:** EMG response of hand and arm muscles (diagram) in response to suprathreshold stimulation of the C7 dorsal root with doublets of different frequencies in Mk-KA. Right: Mean and standard error plots showing amplitude of the 2^nd^ response of the doublet normalized to its 1^st^ response at different delays before (black) and after (cyan) intrathecal gabazine administration at the cervical segments of the spinal cord. After GABA_A_ blockade with gabazine, the normalized amplitude of the second response was reduced within the time course of PAD (*: P<0.001; two-tailed bootstrapping with 147 and 148 points for 10 ms delay, 146 and 143 points for 20 ms delay, 149 and 201 points for 25 ms delay, 147 and 145 points for 33 ms delay, 144 and 136 points for 50 ms delay, 137 and 136 points for 100 ms delay, for pre and post drug conditions, respectively).

We then investigated the main point of the controversy, is PAD excitatory or inhibitory to sensory afferents? According to Hari, Lucas-Osma [18], PAD supports sensory transmission by preventing conduction failure in sensory afferents. If this is correct, then blocking PAD should increase the rate of failure and reduce the size of reflex responses evoked during the time course of PAD elicited from an initial pulse. When stimulating dorsal roots at motor threshold (all-or-none responses) and recording hand muscle EMG (**Fig. 2A**), we found that blocking PAD caused up to 30% increase in failure of the second response to a doublet stimulation. Moreover, increased failure occurred only at inter-pulse intervals that coincided with PAD elicited by the first pulse (25 and 50 ms, but not 100 ms, **Fig. 2B** and **Extended Data Fig. 4A**).

In another experiment, we set stimulation above threshold to test the effect of GABA_A_ blockade on reflex amplitude. GABA_A_ blockers caused about 50% reduction in the normalized amplitude of the second reflex in a doublet in a fashion that matched the time course of PAD (**Fig. 2C** and **Extended Data Fig. 4B**). We further explored this concept by analyzing reflex responses elicited by repeated dorsal root stimulation which are known to be affected by frequency-dependent suppression [30]. We found that also in this case subsequent responses to the first pulse were further suppressed when PAD was blocked (**Extended Data Fig. 4C**), despite that the underlying incoming sensory volleys were not affected by the GABA_A_ blockers (**Extended Data Fig. 2C-D**). Interestingly, even responses evoked by a single pulse at 1-2 Hz were suppressed (**Extended Data Fig. 4D**), suggesting a persistent, tonic facilitatory effect of GABA [18].

These results suggest that GABA has a facilitatory action on sensory afferents. Indeed, blocking GABA should increase general neural excitability, as it is inhibitory in central neurons, demonstrated by higher background spiking in the grey matter **(Extended Data Fig. 3)**. Therefore, the observed reduction in reflex output here indicates that GABA is facilitating sensory inputs through GABA_A_. Importantly, since GABA_A_ is mainly expressed at nodes of Ranvier and not at the terminals [17], our results should be interpreted as axonic nodal mechanisms.

So far, we explored the effects of PAD on spinal sensorimotor processes. However, sensory inputs drive conscious perception of movement as well, and therefore it’s reasonable to ask whether modulation of sensory inputs by spinal PAD affects supraspinal processing of these inputs. Differently from most classical studies on PAD, our in-vivo primate model allowed us to record the propagation of sensory volleys from the spinal cord to the thalamus and into the somatosensory cortex (**Fig. 3A**). Synchronized neural recordings from the thalamus and somatosensory cortex revealed that ascending volleys and field potentials evoked by the 2^nd^ pulse of a doublet were reduced by about 20% after blocking PAD with gabazine administered at the spinal level (**Fig. 3** and **Extended Data Fig. 5**). Furthermore, in two other monkeys, even the responses to a single pulse (at 1 Hz) were suppressed after bicuculline administration (**Extended Data Fig. 6**), again suggesting a persistent tonic GABAergic facilitation of afferents. Taken together, these results indicate that GABA facilitates sensory transmission up the dorsal columns to the brain. Given the importance of this finding we confirmed it also in mice where we could leverage an isolated spinal cord preparation of GAD2//ChR2 mice. Optogenetic activation of GAD2^+^ GABAergic neurons facilitated ascending sensory volleys in the dorsal columns (**Extended Data Fig. 7**) confirming results in monkeys on ascending afferent input.

**Fig. 3.**
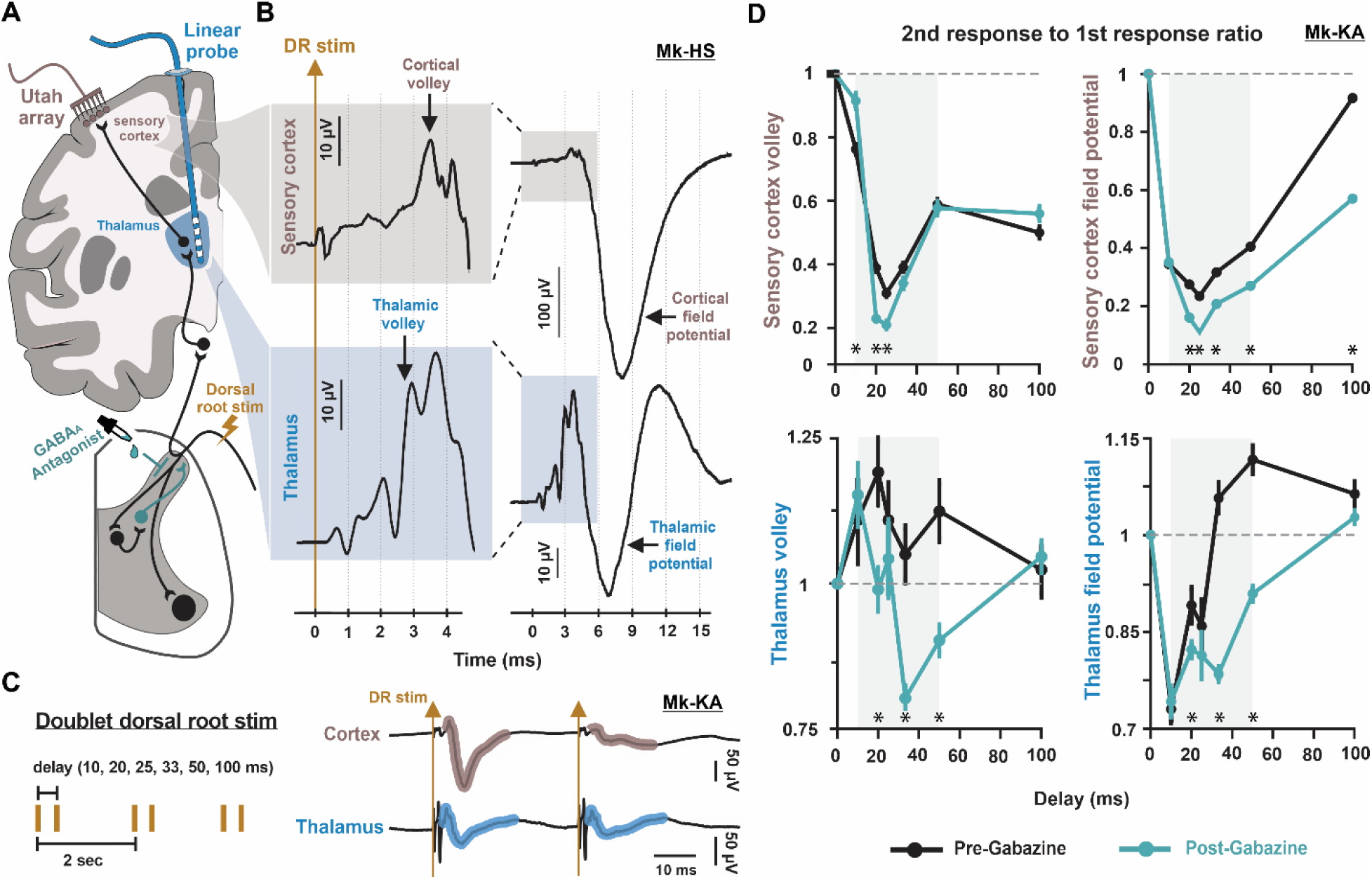
GABA in the spinal cord facilitates transmission of ascending sensory inputs to the cortex. **A**: Schematic diagram showing spinal and supraspinal pathways of primary afferents along with the basic experimental setup. The supraspinal pathway depicted represents the dorsal column-medial lemniscus pathway. **B:** Example traces recorded with the Utah array in the hand area of sensory cortex (top, nude), and a linear probe in thalamus (bottom, blue) showing evoked potentials in both areas in response to a single C6 dorsal root stimulus in Mk-HS. The shaded inset is a magnification of the early parts of the response (see scale) showing the incoming volleys that precede the evoked potentials. **C:** Stimulation protocols of the C7 dorsal root with doublets of different frequencies, and example evoked responses in sensory cortex and thalamus. **D:** Mean and standard error plots summarizing the amplitude of the volley (left) and evoked potential (right) of the 2^nd^ response normalized to its 1^st^ response in the sensory cortex (top) and thalamus (bottom) before (black) and after (cyan) intrathecal gabazine administration at the cervical segments of the spinal cord for Mk-KA (*: P<0.001; two-tailed bootstrapping with 74 and 74 points for 10 ms delay, 72 and 73 points for 20 ms delay, 101 and 75 points for 25 ms delay 73 and 74 points for 33 ms delay, 68 and 72 points for 50 ms delay, 68 and 69 points for 100 ms delay, for pre and post drug conditions, respectively).

In summary, our results help paint a sophisticated picture of sensory modulation. Taken together, they corroborate the findings from recent literature in rodents suggesting that GABA has a facilitatory function on sensory inputs in the cord. Additionally, we show here that GABA (via GABA_A_ receptors) facilitates ascending sensory inputs into supraspinal structures, thus modulating conscious experience of sensory percepts. Since we applied GABA_A_ antagonists locally in the spinal cord, our results suggest that GABAergic modulation in spinal circuits are mediating these effects possibly via preventing failure of action potential propagation at the numerous branching points of sensory afferents [18].

However, we are not suggesting that GABA is only facilitatory nor that presynaptic inhibition does not exist. Instead, we propose that GABA can modulate sensory inputs in two ways: 1) it can facilitate spike propagation in long branching axons of afferents via GABA_A_ receptors, and 2) it inhibits afferent inputs pre-synaptically at the axon terminals [3, 27], possibly through GABA_B_ receptors [18]. This new perspective proposes a fascinating input-biasing system that can provide task-dependent modulation of sensory inputs. In fact, recent studies show that cutaneous and proprioceptive afferents of the arm are modulated differently during extension and flexion tasks in awake monkeys [26, 31]. Furthermore, our proposed perspective provides a new control mechanism for descending control of afferent activity. In this regard, recent evidence suggests that cortico-spinal tracts can modulate primary afferent inputs through PAD [19, 26, 32].

Finally, our data highlights the extent to which PAD impacts the production of movement even at the hand. Indeed, it could have been argued that PAD would not significantly affect hand motoneurons since they evolved a unique dependency on monosynaptic cortico-spinal inputs [33]. Instead, we showed that blocking GABA_A_ receptors substantially impacted hand muscle reflexes causing up to 30% conduction failure to hand motoneurons, suggesting a prominent role of GABA in fine motor control of primates. Our results hold important implications not only in the understanding of motor control and sensory perception, but also call for a critical reexamination of the impact of neurological disorders on GABAergic function in the light of the fact that GABA can facilitate, not only inhibit peripheral sensory inputs.

## METHODS

### In-vivo Monkey Experiments

#### Animals

All procedures in this study were carried out in accordance with the National Institutes of Health Guide for Care and Use of Laboratory Animals. As part of our effort to minimize the number of animals used, the current dataset is derived from experiments that were designed to answer multiple scientific questions [34]. The datasets presented here were collected from 4 macaque monkeys, consisting of 3 adult *Macaca Fascicularis* (MK-OP: age 6, 6kg, female; MK-JC: age 6, 7.5kg, male; MK-KA: age 6, 6.1kg, female) and 1 adult *Macaca Mulatta* (MK-HS: age 7, 12kg, male). The animals were housed in a primate facility at the University of Pittsburgh with ad libitum access to food and water, in addition to daily enrichments (e.g. puzzles and toys). Animals involved in specific experimental procedures are reported in **Extended Data Table 1**.

#### Surgical procedures

The surgical and experimental procedures were reviewed and approved by the University of Pittsburgh Institutional Animal Care and Use Committee (protocol IS00017081) and were performed by Certified neurosurgeons (Drs. Jorge Gonzalez-Martinez, Peter Gerszten, Vahagn Karapetyan, and Daryl Fields, UPMC, Pittsburgh, USA). Prior to the terminal experiments, animals went through one survival procedure to determine trajectories and depth for electrodes that would be implanted in the thalamus during terminal experiments. The details of this procedure have been reported in a previous publication [34].

#### Drugs and chemicals

To test the role of spinal GABA_A_ receptors in sensory processing, we measured sensory-evoked responses before and after the application of GABA_A_ receptor antagonists, either bicuculline or gabazine. The solution (Bicuculline: 436 uM, Gabazine: 436 uM, GABA_A_ antagonists, Sigma Aldrich) was applied intrathecally to the spinal cord after the dura was opened. For both drugs, the powder was first dissolved in DMSO (100 mM, Cambridge Isotope Labs), which does not have a known effect on sensory axons or reflexes but was nonetheless kept at a minimum (final concentration <0.5%). Then, we diluted the solution to a concentration of 16 ug/100 µl in Dulbecco’s Phosphate Buffered Saline (Sigma Aldrich). This concentration was chosen based on studies in rodents [17, 18] with relative scaling based on the size of the monkey spinal cord. During surgery, after all baseline experiments were executed, we first added an initial bolus of 100 µL solution followed by 50 µL or 100 µL every 30 min until the experiments were completed.

#### Terminal procedures

During terminal experiments, anesthesia was induced with ketamine (10 mg/kg, i.m.) and maintained throughout the rest of the experiment with intravenous infusion of propofol (9-27 mg/kg/h) and fentanyl (5-42 µg/kg/h) until animals were euthanized with a single injection of pentobarbital (86 mg/kg, i.v.) after data collection.

#### Muscle and peripheral nerve implantations

We dissected the deep branch of the radial nerve (proprioceptive branch) of the left upper limb and implanted a nerve cuff electrode around it. The nerve branch was verified via electrical stimulation and assessment of muscle responses. We then inserted a pair of needle electrodes into multiple wrist and hand muscles (ECR: extensor carpi radialis, EDC: extensor digitorum communis, FDM: flexor digiti minimi, FCR: flexor carpi radialis, FDC: flexor digitorum communis, APB: abductor pollicis brevis, and biceps) of the same limb.

#### Thalamic depth electrode implantation

We then positioned the animal in a stereotaxic frame (Kopf, Model 1530, Tujunga, CA, USA) in a prone position. The ROSA One(R) Robot Assistance Platform was used to accurately implant a probe in the thalamus contralateral to the muscle and nerve implants [35]. In summary, co-registered MRI and CT images were used to determine a frontal insertion trajectory that kept the motor and somatosensory cortices intact. The robot guided the precise drilling of a penetration hole in the skull and the accurate implantation of a fixation bolt. We then inserted a linear multi-electrode probe through the fixation bolt until it reached the motor-sensory thalamus along the planned trajectory. The correct position of the electrode was confirmed by 1 Hz electrical stimulation of the cervical dorsal roots which evoked volleys and field potentials in the thalamus that preceded those in the sensory cortex (**Fig. 3**).

#### Cortical array implantations

We then implanted an intracortical electrode array in the primary somatosensory area (S1). For this, we performed a craniotomy over the central sulcus and opened the dura to expose cortical tissue contralateral to the hand where EMG was recorded. The position of S1 was determined in relation to the hand representation of the motor cortex which was verified by bipolar surface electrical stimulation and monitoring EMG output in hand and wrist muscles. We then implanted a Utah Array into S1 using a pneumatic impactor system (Blackrock Microsystems).

#### Spinal root and intraspinal implantations

To gain access to spinal sensorimotor circuits for forelimb muscles, we performed a dorsal laminectomy from C3 to T1 vertebrae, and carefully opened the dura. We then placed a bipolar electrode on the C6-C8 dorsal roots for stimulation and verified that the target root evoked responses in hand and wrist muscles. Immediately caudal to the stimulation electrode, a small rootlet was carefully separated, severed distally to the spinal cord, and its proximal end (attached to the cord) was wrapped around a silver hook electrode to record dorsal root potentials (DRPs). We then implanted an intraspinal electrode array into the C6/C7 spinal segment. To do this, we used fine tipped forceps (Dumont #5) to carefully puncture the pia ∼1.2 mm away from the midline and ipsilateral to the hand where EMG was recorded. We then slowly inserted a linear multi-electrode probe into the cord at a 90° angle, and to a depth of 3 mm, using Kopf micromanipulators (Kopf, Model 1760, Tujunga, CA, USA).

#### Electrical stimulation

We used an AM-Systems stimulator (model 2100, Sequim, WA, USA) for radial nerve and dorsal root stimulation. The dorsal root was stimulated using a bipolar electrode (73602-190/10, Ambu Neuroline, Denmark), and the nerve was stimulated through a bipolar cuff electrode (FNC-2000-V-R-A-30, Micro-Leads Neuro, Ann Arbor, MI, USA). Stimuli were delivered as either singles, doublets (at 10ms, 20ms, 25 ms, 33.3ms, 50 ms, or 100 ms latencies) or bursts (burst duration of 1 s at 10 Hz, 50 Hz, or 100 Hz) of biphasic symmetric square pulses. The frequency of stimulation is specified in the figures for each experiment. The stimulation intensity for each animal/target was set slightly higher than the motor threshold (∼1.2xT, visible movement in the muscle or EMG activity). Detailed information on the amplitudes and pulse widths used for stimulation are reported in **Extended Data Table 2**.

#### Electrophysiological recordings

EMG signals were recorded using a pair of needle EMG electrodes (7 and 13 mm, Rhythmlink) inserted about 1 cm apart in the muscle belly. The differential between the two electrodes was then calculated offline. The reference for All EMG recordings was a needle electrode inserted in the lower back or the tail. The intraspinal field potentials and single units were recorded using a linear 32-channel probe (A1x32-15mm-100-177-CM32, NeuroNexus, Ann Arbor, MI, USA). The probe reference was inserted under the skin next to the probe insertion site. Cortical potentials were recorded using a 48-channel Utah Array (400μm pitch, electrode length 1.0mm Blackrock Microsystems, Salt Lake City, UT, USA). The Utah Array reference was inserted between the dura and the skull. Sensory volleys in the thalamus were recorded using a linear 16-channel Dixi electrode (DIXI MicrodeepR SEEG Electrodes). The contacts verified to be in the thalamus (3-4 channels) were used for analysis and referenced to a nearby electrode in the same probe. Dorsal rootlet potential (DRP) was recorded via wrapping the cut end of a dorsal rootlet around a silver wire that has been soaked in bleach to form a silver chloride coat (Ag-AgCl electrode). This electrode was referenced to another silver electrode inserted under the skin next to the recording site. To improve the DRP signal, the cut end of the rootlet and hook electrode were isolated electrically by carefully lifting them away from the cord tissue and submerging them in a small pool of mineral oil. The DRP was amplified by 1000x through an AC Amplifier (model 1700 AM Systems, Sequim, WA, USA) and acquired through the Ripple Neuro Processor (Grapevine Neural Interface 1.14.3) as an analog input at a sampling frequency of 30 kHz. All other electrophysiological signals were amplified through a Ripple Neuro Nano2 headstage (intracortical potentials) or Nano2-HV headstage (thalamus potentials, intraspinal potentials, and EMG), filtered (High-pass: 1.0 Hz, Low-pass: 7.5 kHz) and digitized at 30 kHz using the Ripple Neuro Grapevine and Trellis software (version 1.13 and 1.14).

*Data analysis.* The data was stored on a computer and analyzed offline using Spike2 software (version 10.18, CED, UK) and MATLAB (version R2020b). The amplitude of all signals was measured relative to the pre-stimulus baseline (5-10 ms).

#### Analysis of dorsal root and intraspinal field potentials

The DRP recording from the hook electrode was high-pass filtered at 1 Hz to remove baseline drifting, as well as a 60 Hz notch filter to remove the line noise. We then extracted a window of 160 ms (10 ms pre- and 150 ms post-stimulation). This signal sometimes contained an electrocardiogram (ECG) component. Care was taken to minimize the potential interference from the ECG signal by excluding trials where any of the ECG waveforms overlapped with the post-stimulus analysis window of afferent volley (2-4 ms) or PAD (20-100 ms). Intraspinal array data were bandpass filtered between 1 Hz and 4.9 kHz with a 2nd order Butterworth filter. We extracted the same time window of 160 ms (10 ms pre- and 150 ms post-stimulation) as for the DRP to capture sufficient baseline signal and cover the entirety of the afferent volley and evoked PAD. We quantified the afferent volleys by the peak-to-peak amplitude within the 2-4 ms post-stimulus window, and the PAD by the peak amplitude.

#### Analysis of single unit activity

To extract single unit spiking events, the broadband data was digitally filtered with a high-pass filter of 250 Hz, and spike waveforms (consisting of 52 samples, 1.7 ms) were extracted if they cross a threshold of 3 to 4 standard deviations (SD) of the baseline. The spikes were then sorted with k-means clustering in the Plexon Offline Sorter (Dallas, Texas). Before spike sorting, we concatenated trials of the same stimulation condition before and after bicuculline to ensure we were comparing the changes in firing activity of the same neuron. Single unit activity was presented as stimulation aligned raster plots with the same time window as described above (10 ms pre-stimulation and 150 ms post-stimulation). We then quantified stimulation evoked activity with peri-stimulus triggered histograms (PSTH), defined by average firing probability across all afferent stimulations, calculated with non-overlapping time bins of 2 ms. Units with a distinct peak in PSTH following stimulation were classified as primary afferent related neurons, while other units were classified as spontaneous firing neurons.

#### Analysis of multiunit activity

We computed the spike counts of the extracellular multiunit spiking activity captured by the entire linear probe. To do so, we applied a 3rd order Butterworth digital filter (800-5000 Hz) and detected spikes using threshold crossing, like in the single unit activity analysis. We then removed the stimulation artifact by blanking responses that appeared following the stimulation pulse (blanking window determined by the stimulation pulse width) and 0.5 ms prior to the stimulation pulse. Next, we quantified the spike counts for each channel 100 ms prior to the stimulation pulse and 350 ms after in a 2.5 ms window bin and a moving window of 0.25 ms. Lastly, we smoothed the spike counts with a Gaussian kernel. Prior to computing PCA, we removed noisy repetitions whose SD was 2 times greater than the total SD spike count across all repetitions plus the total mean of the SD across all repetitions for that condition. Subsequently, we z-scored each condition’s spiking activity and computed PCA.

#### Analysis of muscle activity

The EMG signal was calculated differentially between the two needle electrodes in each muscle and was bandpass filtered between 30 and 500 Hz. The monosynaptic EMG response to each dorsal root stimulus was measured as the shortest-latency peak-to-peak compound action potential response. They were measured between 5-20 ms post the stimulation pulse. In the failure rate analysis of doublet stimulations, peak-to-peak responses within 2 standard deviations of baseline noise were considered a failed response. Rate of failure was calculated within every 10 consecutive stimulation pulses. For the response ratio analysis of doublet stimulations, the response ratio was defined as the response amplitude to the second stimulus divided by the response amplitude to the first stimulus.

#### Analysis of thalamus and cortical field potentials

Intracortical array and thalamic electrode data were bandpass filtered between 10 Hz and 4.9 kHz as well as a 60 Hz notch filter. Then, we extracted a window of 25 ms (5 ms pre-stimulation, 20 ms post-stimulation) for analysis. To identify volleys from dorsal rootlet stimulation in the thalamus and cortex, we calculated an approximated time of 2-5 ms based on conduction speed of antidromic volleys in the spinal cord from thalamic stimulation as reported in a previous publication [34]. We measured the thalamic and cortical volleys within the 2-5 ms window, and the evoked field potentials within the 5-20 ms window. The response ratio for doublet stimulations was calculated similarly as for muscle response.

#### Statistical analysis

Statistical comparisons were performed for neural and muscular responses to stimulation before and after bicuculline. The bootstrapping method, a non-parametric test that makes no assumptions on data distribution, was used for all comparisons. Samples of the same size as the input datasets were drawn 10,000 times from each input dataset with replacement, then the differences of means were calculated to generate a distribution to find confidence intervals. We used two-tailed bootstrapping at confidence intervals of 95%, 99%, and 99.9% (alphas of 0.05, 0.01, and 0.001), where the null hypothesis was rejected if 0 was not within the confidence interval.

### Ex-vivo Mouse Experiments

#### Animals

All experimental procedures were approved by the University of Alberta Animal Care and Use Committee, Health Sciences division (ACUC protocol nos. AUP00000224 and AUP00002891) in accordance with Canadian Council on Animal Care guidelines. Adult mice (2.5–6 months old, both female and male equally; strain detailed below) were used in this study. Mice were caged in groups of 2–4 and maintained on a 12-hour light/dark cycle, in stable conditions of temperature and humidity, with food and water ad libitum.

We used a strain of mice with Cre expressed under the endogenous *Gad2* promotor region (*Gad2tm1(cre/ERT2)Zjh* mice, Jackson Laboratory, 010702; CreERT2 fusion protein expressed under control of the endogenous *Gad2* promotor). *Gad2* encodes the enzyme glutamate decarboxylase 2 (also called GAD65), which is unique to axoaxonic contacting GABAergic neurons that project to the VH (details can be found in our previous studies [18]). These GAD2+ neurons were activated optogenetically using ChR2. CreER is an inducible by injecting tamoxifen in adult mice (4–6 weeks old) to prevent developmental issues of earlier induction of Cre and studied >1 month later. Genotyping was performed according to Jackson Laboratory protocols via polymerase chain reaction (PCR) of ear biopsies using specific primers. The data presented in Extended data Fig. 7 were collected using the whole-tissue sacrocaudal portion of the spinal cord. This is the only part of the adult spinal cord that survives whole ex vivo, allowing axon conduction to be assessed along several spinal segments. Furthermore, it has very similar spinal circuitry, reflex and motor neuron properties to those seen in the hindlimb of other preparations.

#### Sacrocaudal spinal cord extraction

Mice were anesthetized with urethane (for mice, 0.11 g/100 g, maximum of 0.065 g); a laminectomy was performed; and then the entire sacrocaudal spinal cord was dissected with all its spinal roots attached and bathed in oxygenated dissection solution (details below). Spinal roots were removed, except the sacral S2, S3, S4 on both sides of the cord. The tissue was allowed to set for 1.5 hours in the dissection chamber (at 20 °C), to wash out the residual anesthetic before recording. The cord was then transferred to a recording chamber perfused with ACSF (recycled in a closed system) at >3 ml/min and maintained at room temperature. The dorsal surface of the cord was oriented upwards, and the laser beam used for optogenetics was focused vertically downward on the GAD2 neurons, as detailed below.

#### Dorsal column stimulation and ascending volley recording

A tungsten electrode was inserted into the dorsal column at the S4 segment to stimulate sensory axons. Stimulation was delivered as constant current short pulses (0.1 ms) at 1.5xT (T, threshold for fast spike initiation) to selectively target proprioceptive afferents. To record the composite intracellular response of many sensory axons, we employed a grease gap method by recording from dorsal roots grease gap methods, where a high impedance seal on the axon allows the recording to reflect IC potentials. The Dorsal roots were cut short and their proximal stump mounted on silver–silver chloride wires close to the dorsal column (2 mm) to record compound action potentials of sensory axons (sensory volleys). The roots were covered with grease (a 3:1 mixture of petroleum jelly and mineral oil) and then surrounded by a more viscous synthetic high vacuum grease to prevent oil leaking. Return and ground silver–silver chloride wires were immersed in the bath.

#### Optogenetic activation of GAD2^+^ neurons

GAD2^+^ neurons were activated using 447-nm D442001FX laser from Laserglow Technologies. Light derived from the laser was focused longitudinally on the left side of the spinal cord (5 ms pulses, λ = 447nm laser, 0.7 mW/mm^2^) at the level of the DH, to target the epicenter of GAD2^+^ neurons, which are dorsally located. This method has been to effectively activate the target neurons [18]. we added silicon carbide powder (9% wt, Tech-Met) to the grease covering the dorsal roots to make it opaque to light and minimize light-induced artifactual current in the recording wire during optogenetic activation of ChR2. Likewise, we covered the ground and reference wires in the bath with a plastic shield to prevent stray light artifacts.

#### Drugs and solutions

The dissection solution used before recording was composed of (in mM) 118 NaCl, 24 NaHCO_3_, 1.5 CaCl_2_, 3 KCl, 5 MgCl_2_, 1.4 NaH_2_PO_4_, 1.3 MgSO_4_, 25 D-glucose and 1 kynurenic acid. The ACSF used during recording was composed of (in mM) 122 NaCl, 24 NaHCO_3_, 2.5 CaCl_2_, 3 KCl, 1 MgCl_2_ and 12 D-glucose. The solutions were saturated with 95% O_2_/5% CO_2_ and maintained at pH 7.4. To prevent glutamatergic circuit activity from interfering with sensory volleys in the dorsal columns, we blocked glutamate receptors by adding the following drugs to the recording ACSF: APV (NMDA receptor antagonist, 50 µM), CNQX (AMPA antagonist, 50 µM). All drugs and chemicals were purchased from Sigma-Aldrich.

#### Statistical analysis

Data were analyzed with Clampfit 10 (Axon Instruments), Excel version 2206 (Microsoft) and SigmaPlot 14.5 (Systat Software). We used a Student’s t-test or one-way ANOVA (as appropriate) to test for statistical differences between two or more independent comparisons, respectively, with a significance level of P < 0.05, all two-sided. Tests were paired for pre-GAD2^+^ and post-GAD2^+^ activation data but otherwise were unpaired. For multiple comparisons with t-tests or ANOVA, we performed a post hoc Bonferroni correction or Tukey test to determine which pairs of measures likely differed.

## ACKNOWLEDGEMENTS

We thank Zimmer Biomet for providing the ROSA robot for our brain surgeries (we declare no conflict of interest). The study was supported by The National Institutes of Health (R01NS131428) of EP, as well as internal funding from the Department of Physical Medicine and Rehabilitation at the University of Pittsburgh to EP and MC. Additional funding was provided by the Walter L. Copeland Fund to EP and MC, and EMG. CJH and AM were supported by The National Institute of Neurological Disorders and Stroke (NS109552). AM was also supported by the Craig H. Neilsen Foundation fellowship (649297). DJB was supported by The Canadian Institutes of Health Research (MOP 14697 and PJT 165823) and the US National Institute of Health (R01NS47567). This investigation also used resources that were supported by the Southwest National Primate Research Center grant P51 OD011133 from the Office of Research Infrastructure Programs, National Institutes of Health and the NIH grant U42 OD120442.

## AUTHOR CONTRIBUTIONS

MC and EP conceived the study and secured funding. AM, LL, JB, JH, KH, DJB, MC, and EP designed the experiments. LL, JB, JH, EG, MC, and EP designed and implemented hardware and software tools for the experiments. LL, JH, JGM, and EP designed the neurosurgical approach for the brain implants, and JGM performed the surgery. LL, JB, JH, PG and MC designed the neurosurgical approach for the spinal implants and PG performed the surgery. VK and DF assisted with all monkeys’ surgeries. AM, LL, JB, JH, EG, AD, MC, and EP performed the experiments and collected all the data in monkeys. KH performed the experiments and collected all the data in mice. AM, LL, JB and KH analyzed the data and generated the figures. DJB, CJH, MC, and EP helped with data interpretation. The manuscript was originally written by MC, AM, EP, and LL and all authors contributed to its revision. MC, EP and CJH supervised the project.

## EXTENDED DATA FIGURES

**Extended Data Fig. 1.**
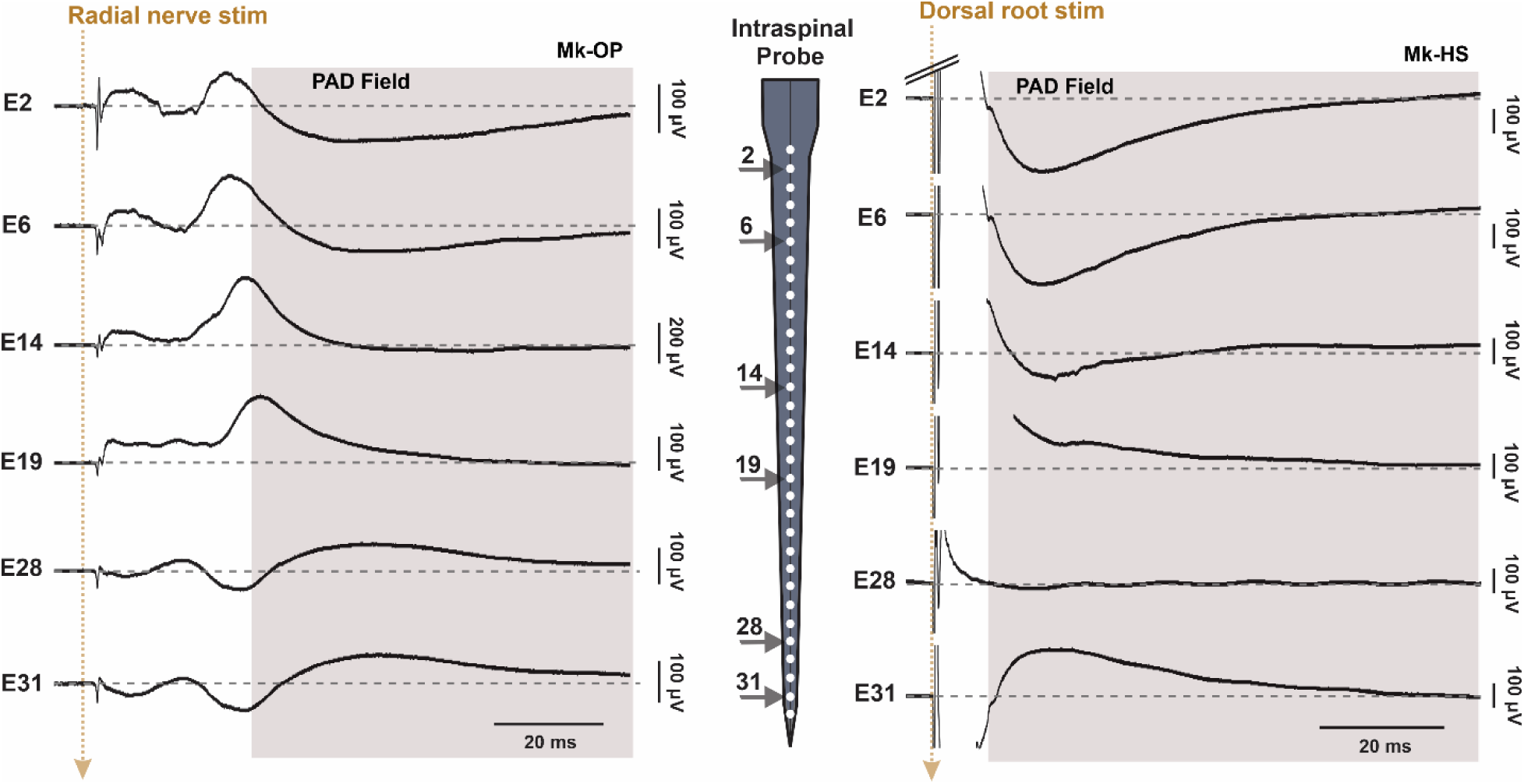
PAD field recording throughout spinal laminae using a multichannel probe. Intraspinal PAD fields recorded using a dorso-ventral linear probe (32-64 channels, schematic in the middle) inserted at the C6-C7 spinal segments in two experiments (same animals in Fig. 1C-D). PAD was evoked by single pulse stimulation (golden dotted arrows, at 1 Hz) of either the deep radial nerve (0.1 ms, **Left**) or the C6 dorsal root (0.4 ms, **Right**) just above the motor threshold. The fields are negative in the dorsal columns and dorsal horn (e.g. E2) and they reverse polarity in the ventral horn (e.g. E31), with some intermediate channels showing no fields (e.g. E19).

**Extended Data Fig. 2.**
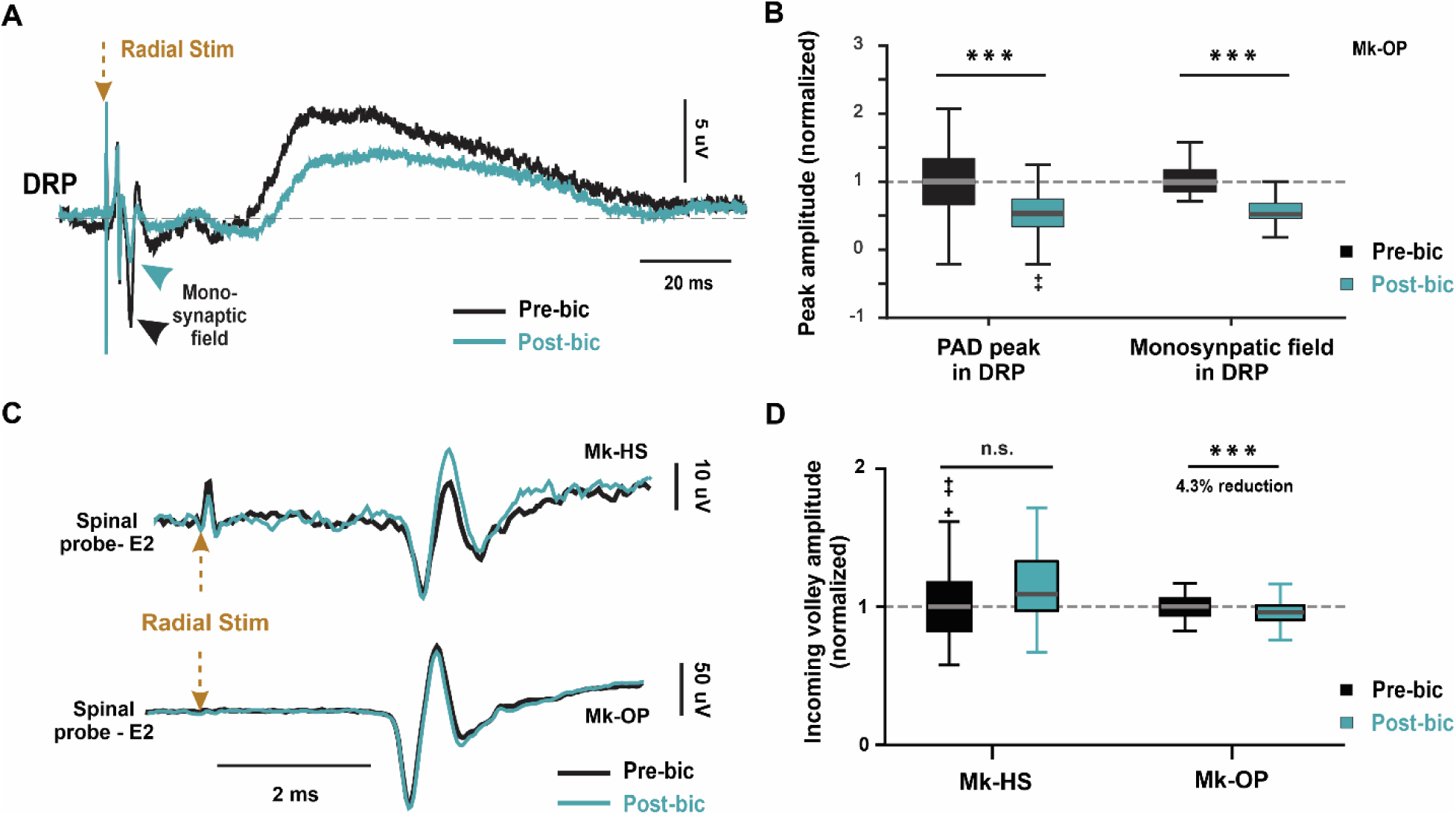
Intrathecal bicuculline administration decreases PAD amplitude in dorsal root potential. **A:** Dorsal root potential (DRP) recorded from a C8 rootlet. The distal end of the rootlet was severed, and the part attached to the cord was wrapped around Ag-AgCl hook electrode for recording. PAD was evoked via single pulse stimulation of the deep radial nerve (1 Hz, 0.1 ms pulse, arrow) before (black) and after (cyan) local intrathecal application of bicuculline on the cervical cord. The arrow heads mark the monosynaptic field evoked by the stimulus. **B:** Bicuculline administration causes reduction in PAD peak amplitude (left) and the evoked monosynaptic field (right) following nerve stimulation (***: P<0.001; two-tailed bootstrapping with 185 and 189 points for pre and post drug conditions, respectively). **C:** Example traces from two experiments showing afferent volleys recorded using one of the electrodes of the intraspinal probe inserted at the cervical segments. The volleys were evoked in response to peripheral stimulation of the deep radial nerve (0.1 ms pulse, at 1 Hz, just above motor threshold) before (Black) and after (Cyan) local intrathecal administration of bicuculline on the cord. **D**: Bicuculline administration causes 16.6% increase in volley amplitude, and yet resulted in > 35% reduction in the evoked PAD (See Fig. 1G). In a different experiment, the afferent volley decreased by only 4.3% after bicuculline, while PAD decreased by 20% in the same experiment (See Fig. 1G). ***: P<0.001; two-tailed bootstrapping with 65 and 61 points for Mk-HS, and 97 and 99 points for Mk-OP. For all boxplots, the whiskers extend to the maximum and minimum, excluding outliers. Central line, top, and bottom of box represent median, 75th, and 25th percentile, respectively.

**Extended Data Fig. 3.**
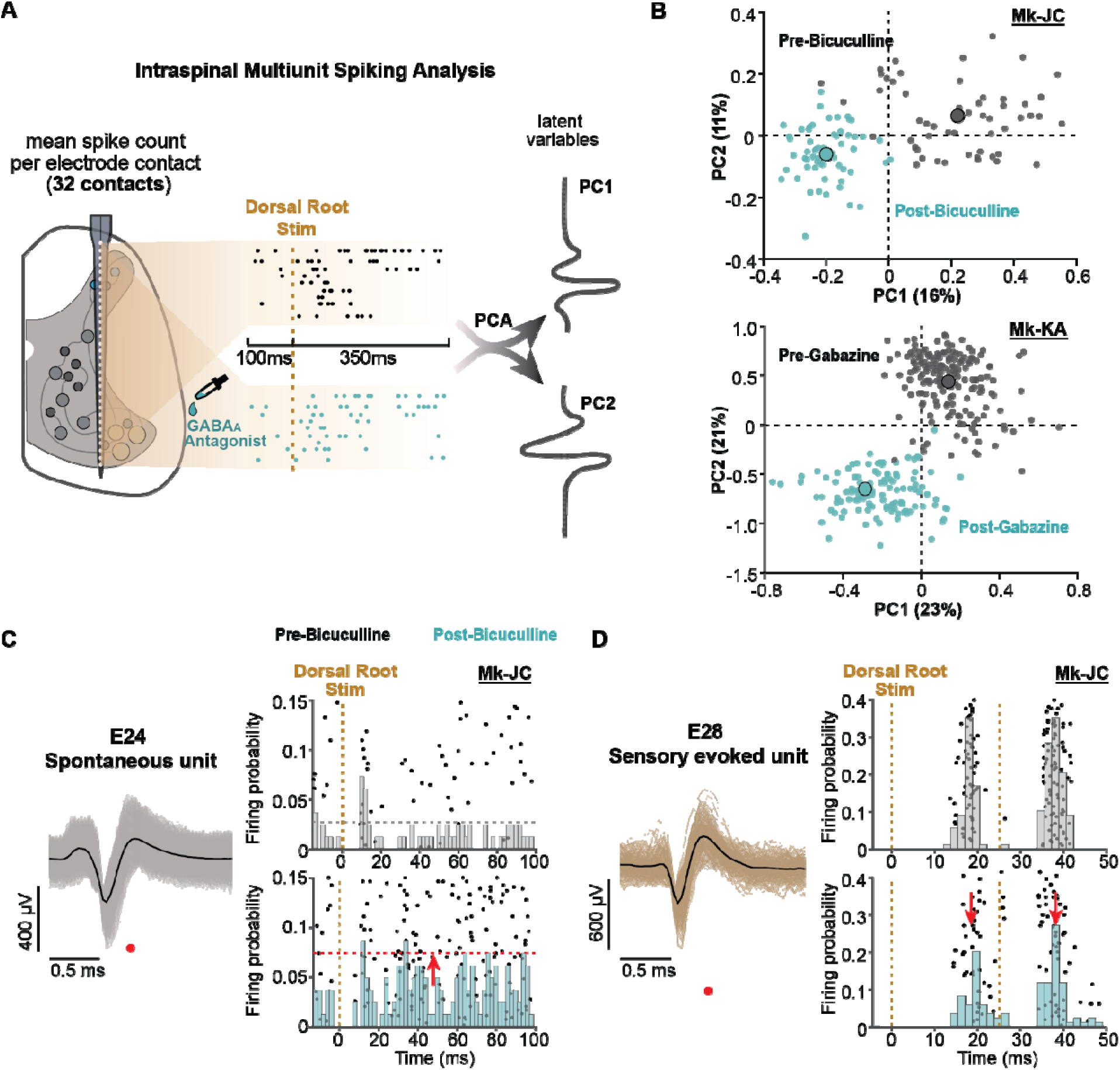
GABA_A_ antagonists increase background neuronal activity in the spinal cord. **A:** Schematic of intraspinal multiunit analysis. Spiking activity across all 32 channels of the linear probe was recorded and aligned by each dorsal root stimulation pulse (100ms pre-stim, and 350ms post-stim), for both before and after GABA_A_ antagonist application. With the mean firing activity of each channel as an input dimension, we performed a principal component analysis (PCA) and extracted the first 2 PCs/latent variables (waveform schematic along probe represent loading coefficient of each channel for the corresponding PC). **B:** Multiunit activity clustered in the PC space before and after bicuculline (top, Mk-JC) and gabazine (bottom, Mk-KA). Intraspinal multiunit activity recorded by the linear probe during single pulse SCS displayed in PC1-2. Each dot represents the mean spike counts across all the dorsoventral channels for each repetition. Centroids represent the mean spike counts across all repetitions. **Top:** Multiunit spiking activity in PC1-2 in Mk-JC (1 Hz, 450 uA SCS; 54 repetitions during control and 60 after the delivery of bicuculline). **Bottom:** Multiunit spiking activity in PC1-2 in Mk-KA (2 Hz, 800 uA SCS; 184 repetitions during control and 118 after the delivery of bicuculline). **C-D:** Example single unit in channel 24 of the spinal probe showing GABA_A_ blocker disinhibited spontaneous firing in the spinal cord (C), while single unit in channel 28 of the same probe shows GABA_A_ blocker reduced sensory input-evoked single unit activity (D).

**Extended Data Fig. 4.**
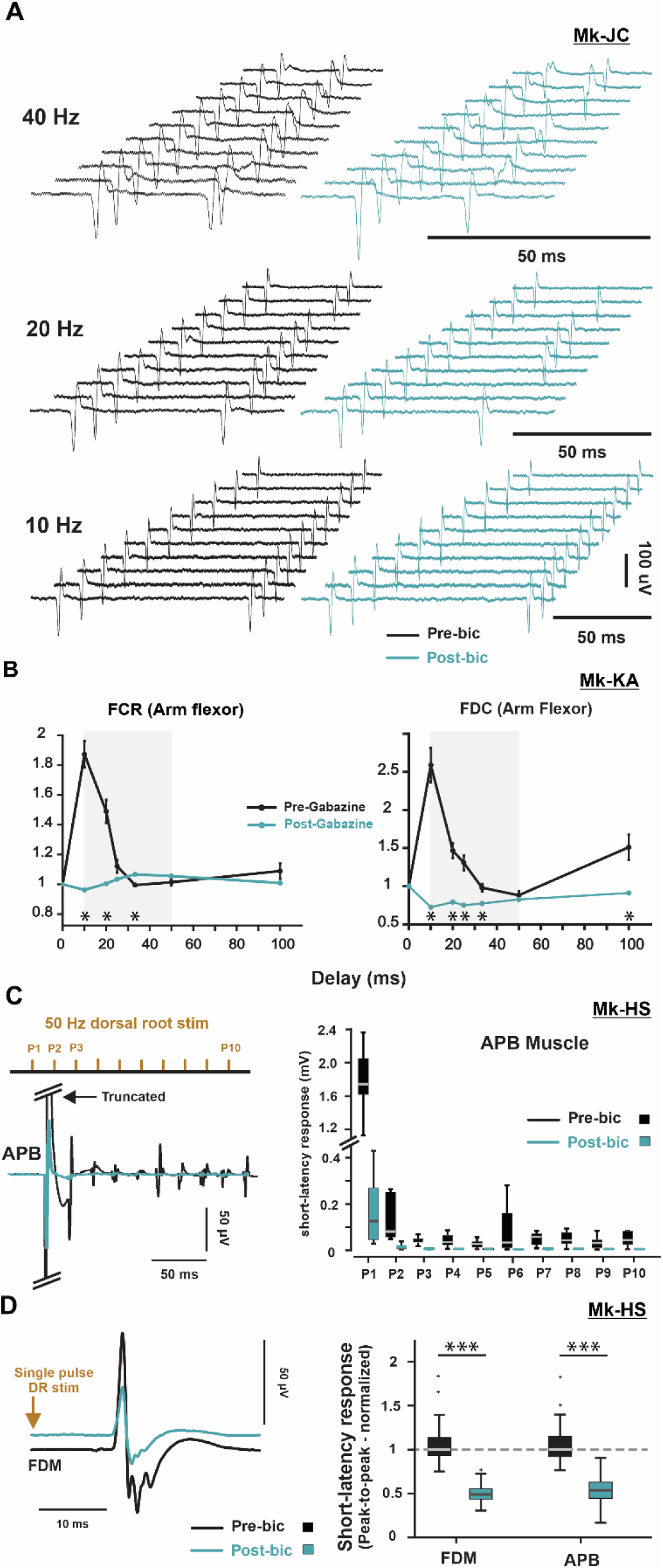
Effect of intrathecal GABA_A_ administration on sensory-evoked motor output. **A**: Raw traces of 10 consecutive EMG responses of the APB muscle to doublet stimulation of the C8 dorsal root at 40 Hz (top), 20 Hz (middle), and 10 Hz (bottom) before (black) and after (cyan) bicuculline local administration to the cervical spinal cord. Note the frequent failure of the second response at 20 and 40 Hz post-bicuculline but none at 10 Hz. The failure rate of the first response was invariably zero at all frequencies pre- and post-bicuculline. **B**: EMG response of FCR and FDC muscles in response to suprathreshold C7 dorsal root stimulation with doublets of different frequencies showing amplitude of the 2^nd^ response of a doublet normalized to its 1^st^ response at different delays before (black) and after (cyan) intrathecal gabazine (GABA_A_ blocker) administration at the cervical segments of the spinal cord. After gabazine, the normalized amplitude of the second response was reduced. *: P<0.001; two-tailed bootstrapping with 147 and 148 points for 10 ms delay, 146 and 143 points for 20 ms delay, 149 and 201 points for 25 ms delay, 147 and 145 points for 33 ms delay, 144 and 136 points for 50 ms delay, 137 and 136 points for 100 ms delay, for pre and post drug conditions, respectively. **C**: EMG response of the APB muscle to a 10-pulses stimulation train to the C6 dorsal root at 50 Hz before (black) and after (cyan) bicuculline in another animal. The amplitude of the short-latency compound action potential was measured at each pulse and plotted (graph on the right). **D**: Bicuculline administration reduces the average EMG response of the FDM and APB muscles evoked by single pulse C6 dorsal root stimulation at 1 Hz. ***: P<0.001; two-tailed bootstrapping with 326 and 201 points for FDM, 358 and 359 points for APB, for pre and post drug conditions, respectively. For all boxplots, the whiskers extend to the maximum and minimum, excluding outliers. Central line, top, and bottom of box represent median, 75th, and 25th percentile, respectively.

**Extended Data Fig. 5.**
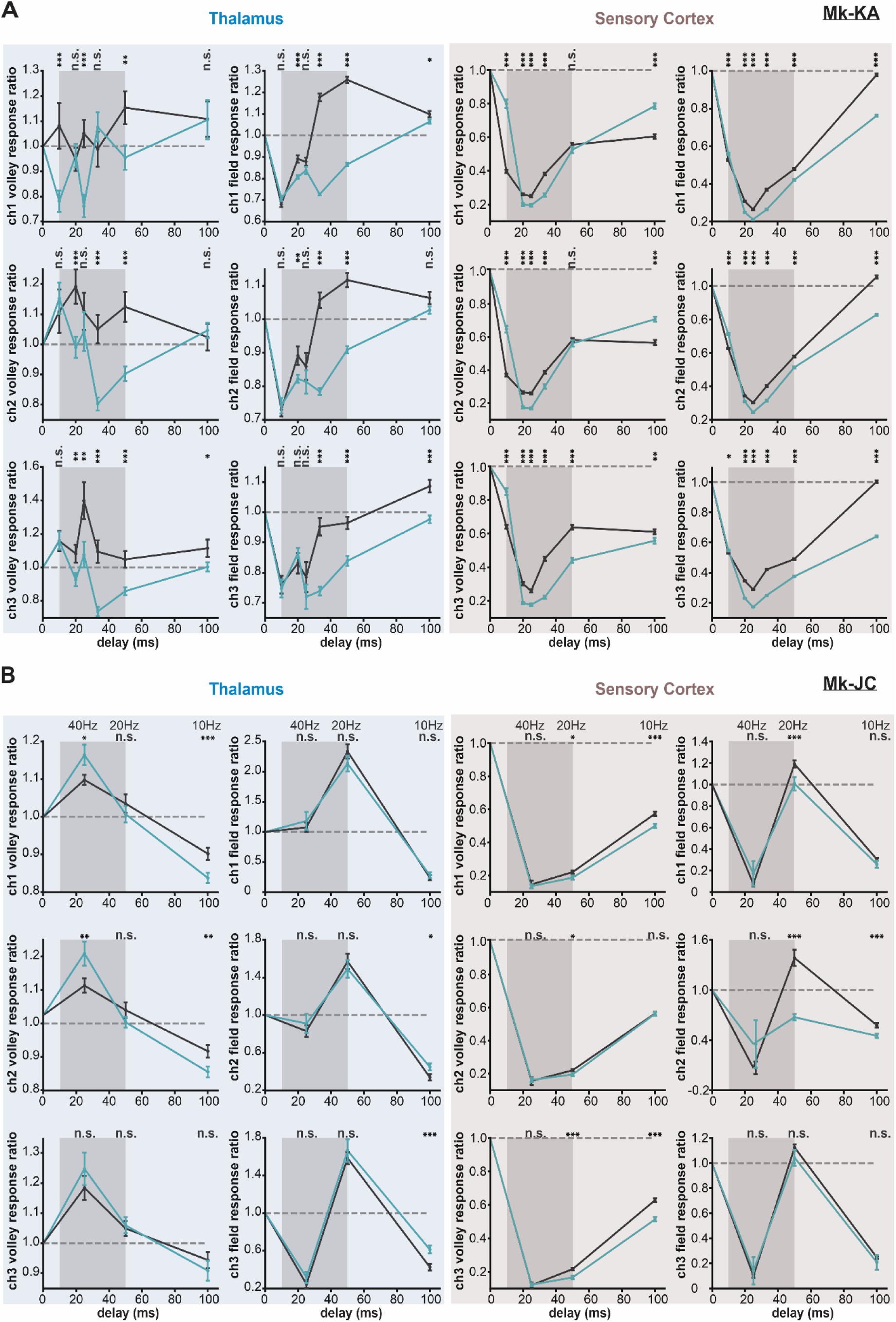
Effect of GABA on ascending afferent inputs. **A:** Mean and standard error plots from more channels of the thalamus probe and S1 Utah array from the same monkey in Fig. 3 (Mk-KA). The graphs show the amplitude of the volley and evoked potential of the 2^nd^ response normalized to its 1^st^ response (doublet stim) in both the thalamus (left) and sensory cortex (right) before (black) and after (cyan) intrathecal gabazine administration at the cervical segments of the spinal cord. *: P<0.05, **: P<0.01, ***: P<0.001; two-tailed bootstrapping with 74 and 74 points for 10 ms delay, 72 and 73 points for 20 ms delay, 101 and 75 points for 25 ms delay, 73 and 74 points for 33 ms delay, 68 and 72 points for 50 ms delay, 68 and 69 points for 100 ms delay, for pre and post drug conditions, respectively. **B:** Mean and standard error plots from thalamus and cortex of another monkey (animal in figure 2B, Mk-JC) showing reduction in cortical responses at 20 Hz. *: P<0.05, **: P<0.01, ***: P<0.001; two-tailed bootstrapping with 44 and 44 points for 25 ms delay, 43 and 43 points for 50 ms delay, 41 and 41 points for 100 ms delay, for pre and post drug conditions, respectively.

**Extended Data Fig. 6.**
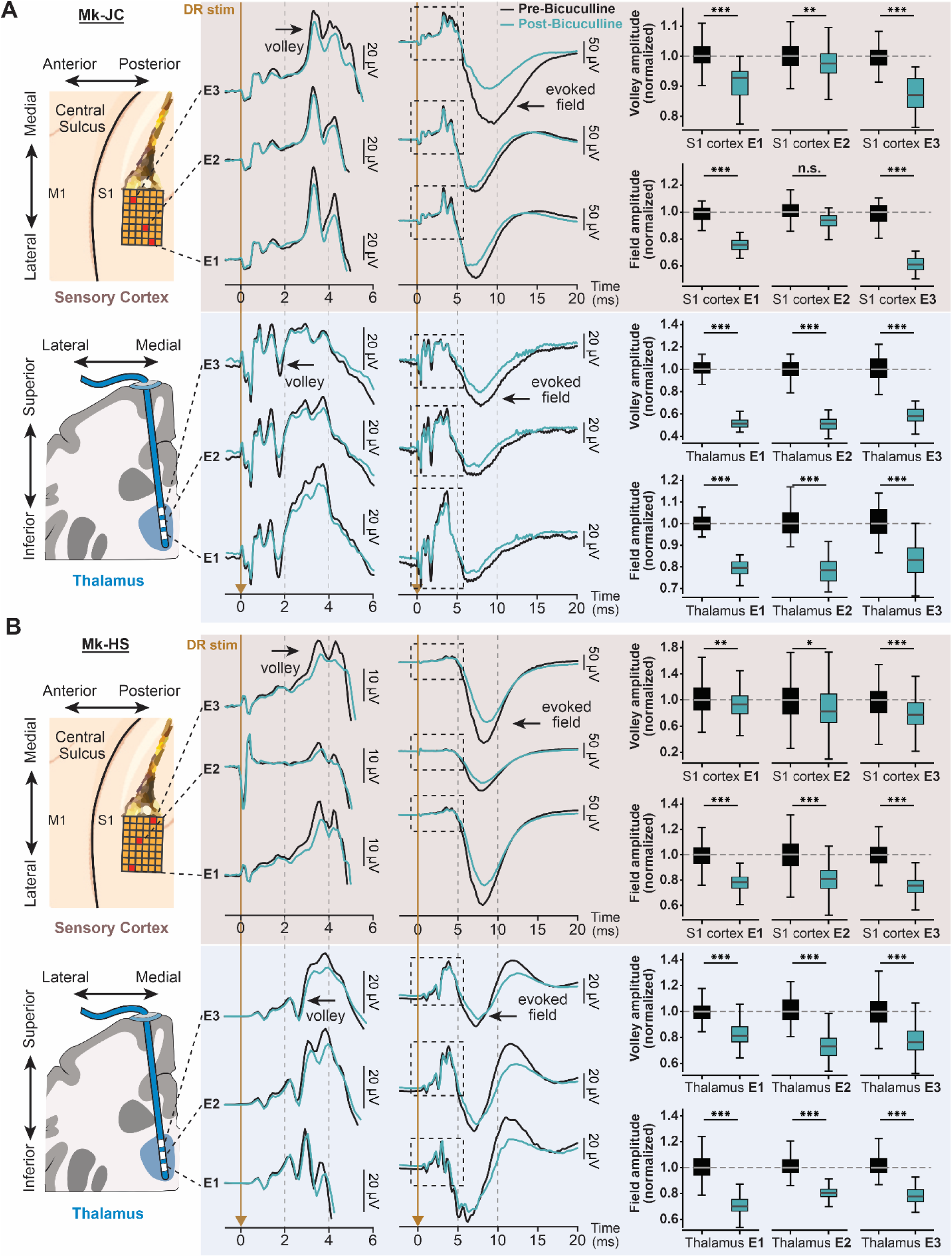
Local bicuculline administration in spinal cord blocks ascending sensory inputs to the thalamus and sensory cortex evoked by a single pulse. **A:** In Mk-JC, **Top left:** average traces of cortical volley and field potential evoked by single pulse DR stimulation at the C7 spinal segment, recorded in 3 channels (E1, E2 & E3) across the Utah array (top, nude). **Bottom left:** average traces of thalamic volley and field potential evoked by the same stimulation pulse, recorded in 3 channels (E1, E2 & E3) of the Dixi probe within the thalamus (bottom, blue). Boxplots quantify the amplitude of the evoked volleys and field potentials normalized to the pre bicuculline condition, showing an overall decrease after bicuculline (*: P<0.05, **: P<0.01, ***: P<0.001; two-tailed bootstrapping with 44 and 44 points for pre and post drug conditions, respectively). **B:** Same as panel A, but in another monkey, Mk-HS, also showing significant decrease of volleys and evoked potentials in thalamus and sensory cortex after bicuculline (*: P<0.05, **: P<0.01, ***: P<0.001; two-tailed bootstrapping with 90 and 90 points for cortical volley, 358 and 359 for cortical evoked potential, and 179 and 180 for thalamus, pre and post drug conditions, respectively). For all boxplots, the whiskers extend to the maximum and minimum, excluding outliers. Central line, top, and bottom of box represent median, 75th, and 25th percentile, respectively.

**Extended Data Fig. 7.**
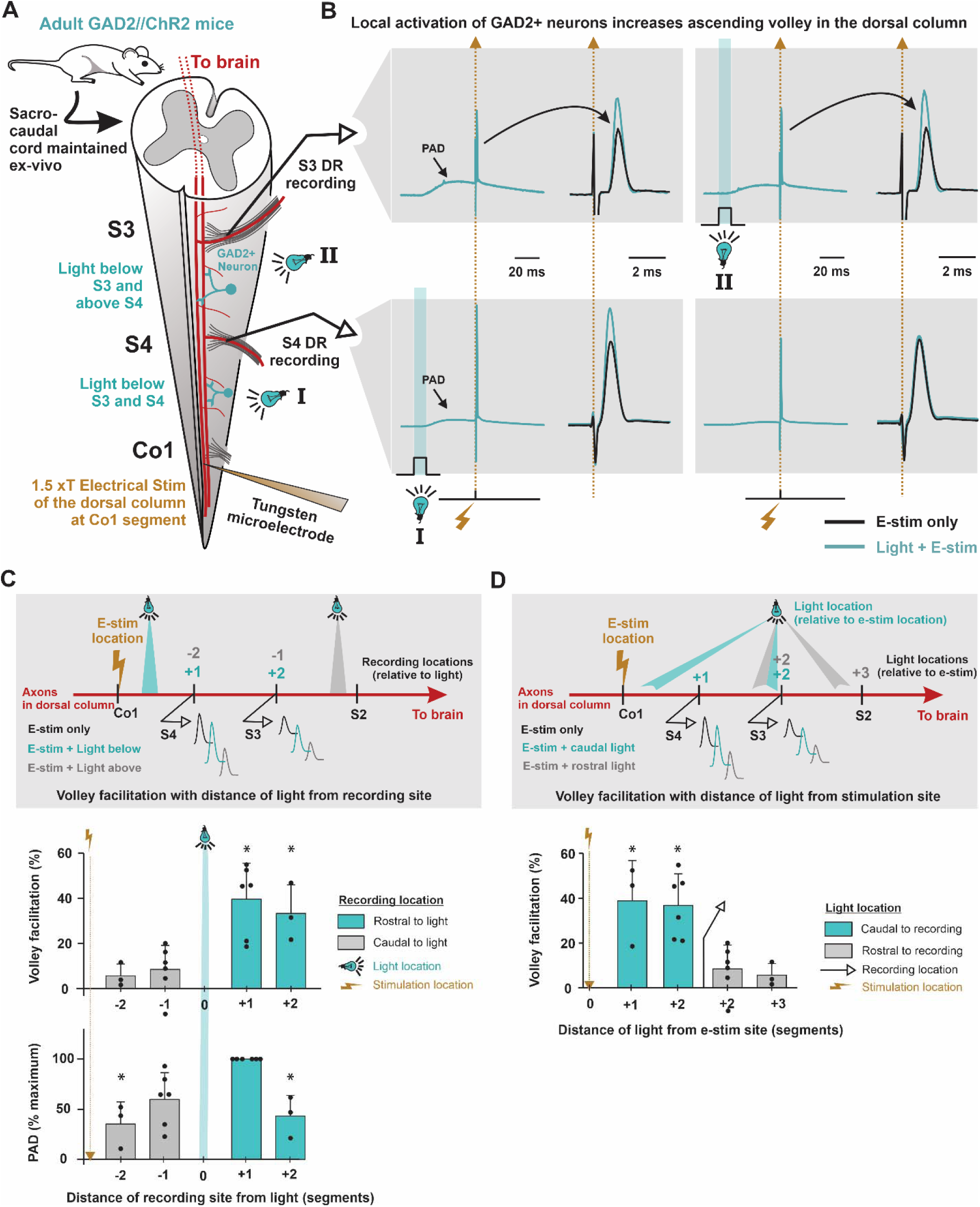
Local optogenetic GABAergic neuron activation increases ascending sensory volley in the large proprioceptive afferents of the dorsal columns. **A:** Schematic of experimental setup. The sacral cord of adult GAD2//ChR2 mice was acutely extracted and maintained ex vivo. The dorsal column at the Co1 coccygeal segment was stimulated extracellularly with a tungsten microelectrode, while recording the ascending sensory volley (compound action potential) to the more rostral segments in the dorsal column ipsilaterally. Action potentials can often fail at the many branch points on the dorsal column axons (proprioceptive Ia afferents’ branch points projects ventrally toward motoneurons), and this would reduce the volley recorded at the surface of the dorsal column. We thus tested this possibility by examining whether optogenetically-evoked PAD through GAD2^+^ neurons could increase the amplitude of the volley recorded at more rostral segments. Electrical stimulation (e-stim) was applied at 1.5xT to activate large myelinated proprioceptive afferents. **B:** Examples of volleys recorded from S4 (bottom) and S3 (top) dorsal roots rostral to the e-stim location. The perfusion solution contained 50 µM of both CNQX and APV to block glutamatergic circuit activity. Amplitude of the volley was facilitated at both locations when GAD2^+^ neurons are optogenetically activated with focal light below S4 (left). As a control, when light was applied just below the S3 root (right), facilitation was seen only at S3 while the S4 volley did not change, confirming that light effects were localized to target segments and not aiding in spike initiation at e-stim site. **C: Top**: Diagram of experimental setup. The volley evoked by e-stim at Co1 was measured at S3 and S4 with light being either caudal or rostral to each root. **Bottom**: Volley facilitation and the amplitude of light-evoked PAD were significantly larger when light is caudal to the recording site (*: P<0.05, significant increase relative to no light condition or light applied rostral to the recording site, n = 6 rootlets recorded from 3 mice). Light applied caudal or rostral to the recording site evoked a PAD visible at the recording site that attenuated with distance. However, the facilitation of the volley by PAD did not attenuate when light was caudal recording site, consistent with PAD facilitating ascending spike conduction along the length of the axon, and not just near the recorded dorsal root filament. **F: Top**: Diagram of experimental setup. The Volley evoked by e-stim at Co1 was measured at S3 and S4 while the light was applied at different distances from the e-stim site. **Bottom**: Similar to the experiments in (C), the volley exhibited more facilitation when light was caudal to the recording electrode than when light was rostral to it (*: P<0.05, significant increase relative to no light condition, n = 6 rootlets from 3 mice). However, this graph clearly shows that the light did not just facilitate the initiation of spikes at the stimulation site, but rather the propagation of the action potential through the axon on route to each recording site.

**Extended Data Table 1.**

| Animal ID | Species | Sex | Age | Body Weight | Experiments |
| --- | --- | --- | --- | --- | --- |
| MK-OP | Macaca Fascicularis | F | 6 | 6 kg | • PAD evoked by nerve stimulation.<br>• PAD fields recorded with intraspinal electrode array.<br>• Suppression of PAD by bicuculline. |
| MK-HS | Macaca Mulatta | M | 7 | 12 kg | • PAD evoked by nerve stimulation.<br>• PAD evoked by dorsal root stimulation.<br>• PAD fields recorded with intraspinal electrode array.<br>• Suppression of PAD by bicuculline.<br>• Suppression of motor responses (EMG) after bicuculline (single pulse and burst stimulation).<br>• Suppression of sensory-evoked potentials in the cortex after bicuculline (single pulse stimulation). |
| MK-JC | Macaca Fascicularis | M | 6 | 7.5 kg | • PAD evoked by dorsal root stimulation.<br>• PAD fields recorded with intraspinal electrode array.<br>• Suppression of PAD by bicuculline.<br>• Suppression of motor responses (EMG) after bicuculline (doublet stimulation).<br>• Suppression of sensory-evoked potentials<br>in the cortex after bicuculline (single pulse and doublet stimulation). |
| MK-KA | Macaca Fascicularis | F | 6 | 6.1 kg | • PAD evoked by dorsal root stimulation.<br>• PAD fields recorded with intraspinal electrode array.<br>• Suppression of PAD by gabazine.<br>• Suppression of motor responses (EMG) after gabazine (doublet stimulation).<br>• Suppression of sensory-evoked potentials in the cortex after gabazine (doublet stimulation) |

**Extended Data Table 2.**

| Animal ID | Deep Radial Nerve<br>(amplitude, pulse width) | Dorsal Root<br>(amplitude, pulse width, spine level) |
| --- | --- | --- |
| MK-OP | 120μA, 100μs | 1.8mA, 200μs, T1 |
| MK-HS | 100μA, 100μs | 400μA, 200μs, C6 |
| MK-JC | 120μA, 100μs | 400μA, 400μs, C8 |
| MK-KA | 85μA, 50μs | 800μA, 200μs, C7 |

## REFERENCES

1. Matthews, P.B., Muscle Spindles and Their Motor Control. Physiol Rev, 1964. 44: p. 219–88.

2. Ishizuka, N., et al., Trajectory of group Ia afferent fibers stained with horseradish peroxidase in the lumbosacral spinal cord of the cat: three dimensional reconstructions from serial sections. J Comp Neurol, 1979. 186(2): p. 189–211.

3. Fink, A.J., et al., Presynaptic inhibition of spinal sensory feedback ensures smooth movement. Nature, 2014. 509(7498): p. 43–8.

4. Eccles, J.C., R.M. Eccles, and F. Magni, Central inhibitory action attributable to presynaptic depolarization produced by muscle afferent volleys. J Physiol, 1961. 159(1): p. 147–66.

5. Gotch, F. and V.A.H. Horsley, *VI.* Croonian Lecture.—On the mammalian nervous system, its functions, and their localisation determined by an electrical method. Philosophical Transactions of the Royal Society of London. , 1891(182): p. 267–526.

6. Eccles, J.C., R.F. Schmidt, and W.D. Willis, Presynaptic inhibition of the spinal monosynaptic reflex pathway. J Physiol, 1962. 161(2): p. 282–97.

7. Eccles, J.C., *Postsynaptic and Presynaptic Inhibitory Actions in the Spinal Cord.* Progress in Brain Research, Elsevier, 1963. 1: p. 1–22.

8. Eccles, J.C., R. Schmidt, and W.D. Willis, Pharmacological Studies on Presynaptic Inhibition. J Physiol, 1963. 168(3): p. 500–30.

9. Gray, E.G., A morphological basis for pre-synaptic inhibition? Nature, 1962. 193: p. 82–3.

10. Betley, J.N., et al., Stringent specificity in the construction of a GABAergic presynaptic inhibitory circuit. Cell, 2009. 139(1): p. 161–74.

11. Sung, K.W., et al., Abnormal GABAA receptor-mediated currents in dorsal root ganglion neurons isolated from Na-K-2Cl cotransporter null mice. J Neurosci, 2000. 20(20): p. 7531–8.

12. Gallagher, J.P., H. Higashi, and S. Nishi, Characterization and ionic basis of GABA-induced depolarizations recorded in vitro from cat primary afferent neurones. J Physiol, 1978. 275: p. 263–82.

13. Alvarez-Leefmans, F.J., et al., Intracellular chloride regulation in amphibian dorsal root ganglion neurones studied with ion-selective microelectrodes. J Physiol, 1988. 406: p. 225–46.

14. Lamotte D’Incamps, B., et al., Reduction of presynaptic action potentials by PAD: model and experimental study. J Comput Neurosci, 1998. 5(2): p. 141–56.

15. Rudomin, P. and R.F. Schmidt, Presynaptic inhibition in the vertebrate spinal cord revisited. Exp Brain Res, 1999. 129(1): p. 1–37.

16. Engelman, H.S. and A.B. MacDermott, Presynaptic ionotropic receptors and control of transmitter release. Nat Rev Neurosci, 2004. 5(2): p. 135–45.

17. Lucas-Osma, A.M., et al., Extrasynaptic alpha(5)GABA(A) receptors on proprioceptive afferents produce a tonic depolarization that modulates sodium channel function in the rat spinal cord. J Neurophysiol, 2018. 120(6): p. 2953–2974.

18. Hari, K., et al., GABA facilitates spike propagation through branch points of sensory axons in the spinal cord. Nat Neurosci, 2022. 25(10): p. 1288–1299.

19. Metz, K., et al., Facilitation of sensory transmission to motoneurons during cortical or sensory-evoked primary afferent depolarization (PAD) in humans. J Physiol, 2023. 601(10): p. 1897–1924.

20. Metz, K., et al., Post-activation depression from primary afferent depolarization (PAD) produces extensor H-reflex suppression following flexor afferent conditioning. J Physiol, 2023. 601(10): p. 1925–1956.

21. Szabadics, J., et al., Excitatory effect of GABAergic axo-axonic cells in cortical microcircuits. Science, 2006. 311(5758): p. 233–5.

22. Zorrilla de San Martin, J., F.F. Trigo, and S.Y. Kawaguchi, Axonal GABA(A) receptors depolarize presynaptic terminals and facilitate transmitter release in cerebellar Purkinje cells. J Physiol, 2017. 595(24): p. 7477–7493.

23. Howell, R.D. and J.R. Pugh, Biphasic modulation of parallel fibre synaptic transmission by co-activation of presynaptic GABAA and GABAB receptors in mice. J Physiol, 2016. 594(13): p. 3651–66.

24. Weiler, J., P.L. Gribble, and J.A. Pruszynski, Spinal stretch reflexes support efficient control of reaching. J Neurophysiol, 2021. 125(4): p. 1339–1347.

25. Weiler, J., P.L. Gribble, and J.A. Pruszynski, Spinal stretch reflexes support efficient hand control. Nat Neurosci, 2019. 22(4): p. 529–533.

26. Tomatsu, S., et al., Presynaptic gating of monkey proprioceptive signals for proper motor action. Nat Commun, 2023. 14(1): p. 6537.

27. Seki, K., S.I. Perlmutter, and E.E. Fetz, Sensory input to primate spinal cord is presynaptically inhibited during voluntary movement. Nat Neurosci, 2003. 6(12): p. 1309–16.

28. Lloyd, D.P. and I.A. Mc, On the origins of dorsal root potentials. J Gen Physiol, 1949. 32(4): p. 409–43.

29. Barron, D.H. and B.H. Matthews, The interpretation of potential changes in the spinal cord. J Physiol, 1938. 92(3): p. 276–321.

30. Mahrous, A., et al., Muscle spasms after spinal cord injury stem from changes in motoneuron excitability and synaptic inhibition, not synaptic excitation. J Neurosci, 2023.

31. Confais, J., et al., Nerve-Specific Input Modulation to Spinal Neurons during a Motor Task in the Monkey. J Neurosci, 2017. 37(10): p. 2612–2626.

32. Dallmann, C.J., et al., Presynaptic inhibition selectively suppresses leg proprioception in behaving Drosophila. bioRxiv, 2023.

33. Lemon, R.N., Descending pathways in motor control. Annu Rev Neurosci, 2008. 31: p. 195–218.

34. Ho, J.C., et al., Targeted Deep Brain Stimulation of the Motor Thalamus Facilitates Voluntary Motor Control after Cortico-Spinal Tract Lesions. medRxiv, 2023.

35. Ho, J.C., et al., Robot Assisted Neurosurgery for High-Accuracy, Minimally-Invasive Deep Brain Electrophysiology in Monkeys. Annu Int Conf IEEE Eng Med Biol Soc, 2022. 2022: p. 3115–3118.

